# Single cell mapping of the human endometrium and first trimester decidua identifies distinct epithelial and stromal cell contributions to fertility

**DOI:** 10.1101/2025.04.24.650488

**Authors:** Gregory W. Burns, Emmanuel N. Paul, Manisha Persaud, Qingshi Zhao, Rong Li, Kristin Blackledge, Jessica Garcia de Paredes, Pratibha Shukla, Ripla Arora, Anat Chemerinski, Nataki C. Douglas

## Abstract

The human endometrium undergoes dynamic changes across the menstrual cycle to establish a receptive state for embryo implantation. Using bulk and single-cell RNA sequencing, we characterized gene expression dynamics in the cycling endometrium and the decidua from early pregnancy. We are the first to demonstrate that during the mid-secretory phase—the period encompassing the window of implantation—secretory glandular epithelial cells undergo notable transcriptional changes in cell-cell communication. Through comprehensive analyses, we identified the *Glandular Epithelium Receptivity Module* (GERM) signature, comprising 556 genes associated with endometrial receptivity. This GERM signature was consistently perturbed across multiple datasets of endometrial samples from women with impaired fertility, validating its relevance as a marker of receptivity. In addition to epithelial changes, we observed shifts in stromal cell populations, notably involving decidual and senescent subsets, which also play key roles in modulating implantation. Together, these findings provide a high-resolution transcriptomic atlas of the receptive and early pregnant endometrium and shed light on key molecular pathways underlying successful implantation.

## Introduction

Only 30-40% of ovulatory human menstrual cycles result in spontaneous pregnancy^1^. Coordinated actions of ovarian-derived estradiol and progesterone orchestrate changes in the cellular compartments of the endometrium, including the stromal, epithelial, immune, and endothelial cells^2^, to prepare the endometrium for pregnancy. During the follicular phase of the menstrual cycle, a dominant follicle secretes increasing amounts of estradiol, eventually reaching the threshold required to initiate the luteinizing hormone (LH) surge and trigger ovulation^3–5^. Following ovulation, progesterone secreted from the corpus luteum exerts its actions on the endometrium resulting in a differentiated tissue capable of embryo implantation^6^. A brief period of endometrial receptivity follows; it is aptly termed the window of implantation (WOI) and occurs during the mid-secretory phase between 6 and 10 days after the LH surge^7^. This temporally restricted window is characterized by global changes in gene expression, and an increase in secretory and metabolic activity^8^. Attachment of the blastocyst stage embryo to the luminal epithelium is followed by a cascade of events including embryo-derived trophoblast cell invasion of the endometrial stroma and spiral arterioles^9^. Thus, pregnancy initiation depends on the successful crosstalk between a competent embryo and a receptive endometrium, which is defined by the cellular, molecular and structural milieu. Abnormalities in endometrial or embryonal development, or dyssynchrony between these two elements, could result in implantation failure or pregnancy loss early in gestation^10^.

Assisted reproductive technology (ART) is commonly used to overcome infertility. With *in vitro* fertilization (IVF) and preimplantation genetic testing, embryos can be assessed for quality and ploidy to identify those with the highest chance of success. Studies comparing the endometrium of women in natural menstrual cycles and those undergoing ovarian stimulation have sought to identify factors associated with an implantation-competent secretory phase endometrium^11–13^. Ovarian stimulation induces a dyssynchrony between endometrial glands and stroma, high levels of differentially expressed genes, and significant changes in the immune cell compartment^13^. For this reason, embryo transfer in ART cycles is most often performed separately from ovarian stimulation^14^. However, despite advances in ART, implantation rates still only approach 70%^15^. Additionally, recurrent implantation failure (RIF), defined as failure to achieve pregnancy after several embryo transfers, impacts up to 10% of individuals undergoing IVF and embryo transfer^16,17^. Failed implantation suggests that the optimal endometrial environment has not been fully elucidated, underscoring the need for improved methods to assess and identify an endometrium receptive to embryo implantation.

Current methods to clinically assess endometrial receptivity are limited. In practice, transvaginal ultrasound evaluation of endometrial thickness is the widely accepted approach. In ART cycles, implantation rates are lower when endometrial thickness is < 7 mm^18^, however, the cellular, molecular and structural changes that drive endometrial thickness and define a receptive endometrium are unknown. To address this, endometrial sampling in non-conception cycles, followed by histologic and/or transcriptomic analysis has been explored^19^. It has been proposed that the WOI may have a mRNA signature that can be assessed using transcriptomics technologies. The Endometrial Receptivity Array (ERA) was a tool developed to determine the optimal timing for embryo transfer based on the expression of 238 specific receptivity-associated genes^20^ in an endometrial biopsy sample obtained during a mock, non-conception cycle. The ERA results, which describe the endometrium as pre-receptive, receptive, or post-receptive, could require altering the timing of the subsequent embryo transfer to align with the most receptive day in the cycle. And thus, identification of the gene signature associated with a receptive endometrium, in conjunction with transvaginal ultrasound, would improve pregnancy rates. However, this tool has not been found to improve pregnancy rates for most groups, and recent RCTs evaluating the effectiveness of the ERA have yielded conflicting results^21–23^.

Several gaps remain in our understanding of what constitutes an implantation-competent endometrium. Firstly, it is unclear how gene expression varies throughout the secretory phase or from cycle to cycle in one individual. In addition to the absolute levels of gene expression, capturing the directionality of changes in gene expression is important. Secondly, bulk RNA-sequencing (RNA-seq) analysis of human endometrial biopsies has presented inconsistent results across different studies. Bulk RNA-seq also precludes cell-type specific analysis of gene expression. With the advancement of single-cell RNA-seq (scRNA-seq), it is becoming increasingly clear that various endometrial cell types exhibit distinct transcriptomic patterns^24–26^. Therefore, the individual cell types and their transcriptomic changes may both play important roles in endometrial receptivity in the secretory phase. Finally, the ERA was developed on a non-Hispanic white population. The broad applicability of the ERA to women from different race/ethnicities is lacking. Thus, an improved understanding of the changes in the cycling endometrium is critical and will provide the foundation for developing a clinical tool that could accurately predict endometrial receptivity or preparedness and reduce the chance of implantation failure.

To address these gaps, we conducted bulk mRNA-seq and scRNA-seq analysis of endometrial samples obtained from healthy Black and Hispanic women of proven fertility across the menstrual cycle, and from first trimester decidual tissue. Our goal was to better understand stromal and epithelial cell changes that underlie endometrial differentiation in preparation for embryo implantation, with a focus on acquisition of endometrial receptivity and the processes of decidualization and cellular senescence. Herein, we identify a critical role for the secretory glandular epithelium in supporting endometrial receptivity during the window of implantation. To our knowledge, this is the first report establishing this finding in humans, although the requirement of functional glandular epithelium for fertility is well documented in mice and domestic animals^27–31^. Moreover, we underscore the significance of this cell type in fertility across diverse populations.

## Results

### Experimental study design

The endometrium undergoes significant cellular changes across the menstrual cycle, particularly during the secretory phase which follows ovulation and prepares the endometrium for embryo implantation. With pregnancy, the post-implantation endometrium, is referred to as the endometrial decidua^32^. To understand temporal changes in the transcriptomic signature within endometrial cell populations, we enrolled women with regular, ovulatory cycles and proven fertility and those in the first trimester of pregnancy (**Figure 1A**). After enrollment, 30 subjects were assigned to one of four groups that corresponded to endometrial biopsy collection in the proliferative (n = 6), early- (n = 10), mid- (n = 8), or late-secretory (n = 3) phase of the menstrual cycle or for the collection of endometrial deciduae at the time of an elective termination of pregnancy (n=3) (**Table S1**). Study participants were similar with respect to age, BMI, gravidity (number of prior pregnancies), and parity (number of live births) (**Table S1**). 87% of subjects were Black (26/30) and 13% (4/30) were Hispanic. Subjects in the study had a mean (SD) age of 30.2 (5.2) years and a mean (SD) BMI of 33.0 (8.4) kg/m^2^. The median [IQR] number of pregnancies was 3 [2,4] with 2 [2,3] live births. The estradiol level at the time of the biopsy decreased from 160.9 (74.3) pg/mL in the early secretory phase, to 169.0 (119.5) pg/mL in the mid-secretory phase and 120.7 (12.3) pg/mL by the late secretory phase. As expected, the progesterone level was 1.1 (0.8) ng/mL in the early secretory phase, peaked at 11.5 (3.7) ng/mL in the mid-secretory phase, and decreased to 5.3 (0.9) ng/mL in the late secretory phase.

**Figure 1:**
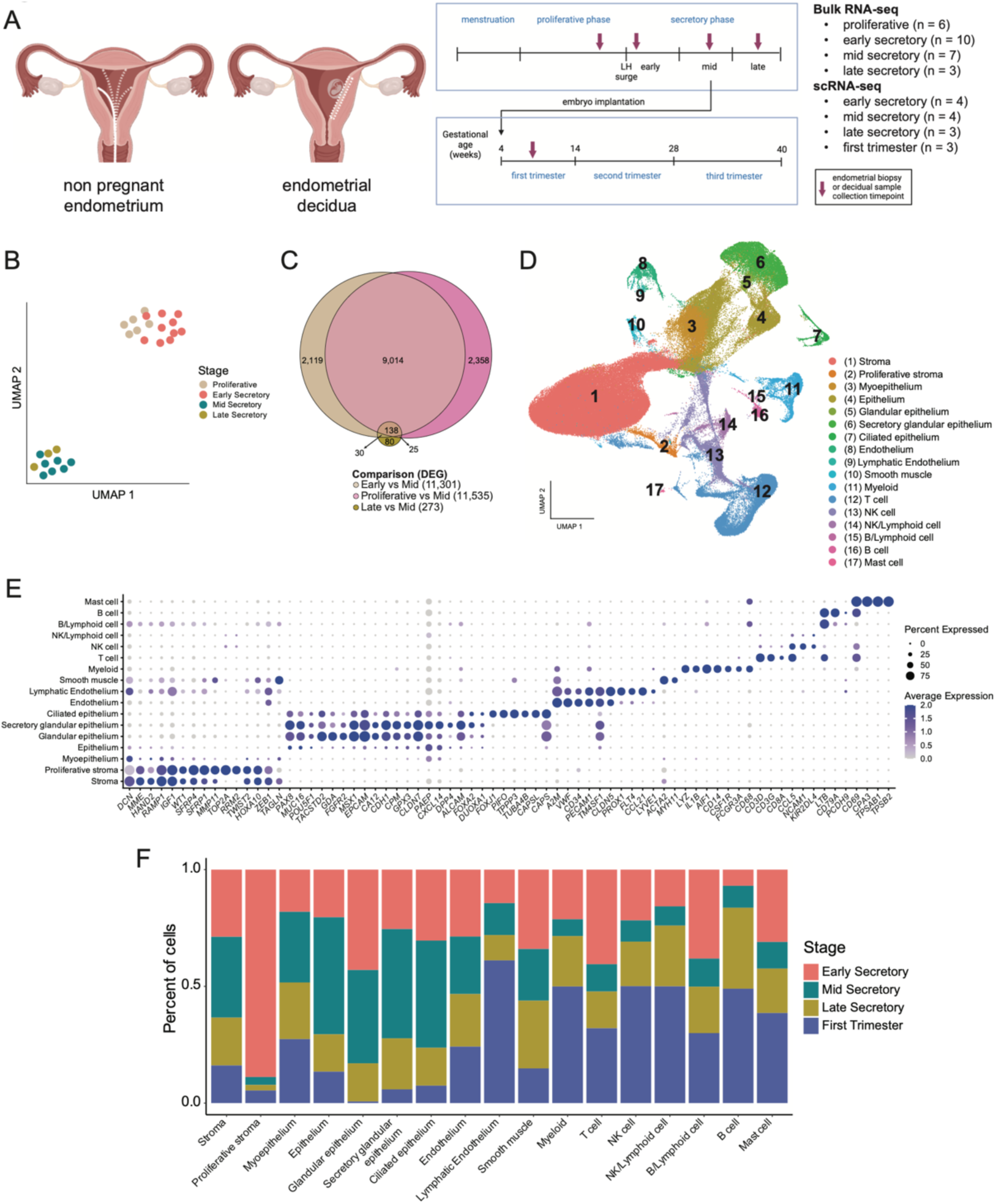
Identification of the transcriptomic profile and dynamic cell populations in the endometrium over time. (A) Summary of sample collection (B) Uniform manifold approximation and projection (UMAP) visualization of RNA-sequencing from proliferative (n = 6), early (n = 10), mid- (n = 7) and late (n = 3) secretory phase human endometrial samples. (C) Euler plot of the differentially expressed genes (DEG) from proliferative vs mid-secretory, early vs mid-secretory, and late vs mid- secretory phase endometrial samples. (D) UMAP visualization of 171,261 isolated cells from human endometrial samples (n = 14). Each cluster (n = 17) represents a cell population with a similar transcriptomic profile. (E) Dotplot for cluster identification using specific markers for cell types from the endometrium. Average gene expression and percentage of cells expressing the specific gene in each cell cluster are shown by the color intensity and the diameter of the dot, respectively. (F) Stacked bar plot showing the proportion of cells in each cluster by stage.

### Bulk and single cell-RNA atlas

To identify global changes in the endometrial transcriptome across the menstrual cycle, bulk mRNA-seq was performed on 26 endometrial biopsy samples from the proliferative and secretory phases. We confirmed menstrual stage separation of the bulk samples with the endest package^33^ for molecular staging of human endometrium (Figure S1A). Uniform Manifold Approximation and Projection (UMAP) resolved two clusters, one cluster containing the proliferative and early secretory phase samples and a second cluster containing the mid- and late secretory phase samples (**Figure 1B**), suggesting a separation in the transcriptomic profiles of these endometrial samples into two groups. This observation was supported by the number of differentially expressed genes (DEG) found between proliferative and early secretory phase samples (n = 205 DEG) compared to proliferative versus mid-secretory (n = 11,535 DEG); early versus mid-secretory (n = 11,301 DEG); and late versus mid-secretory (n = 273 DEG) phase samples (**Figure 1C**, **Tables S2-5**). Moreover, over 9,000 DEG were in common between the proliferative versus mid-secretory or early versus mid-secretory phase samples.

We used scRNA-seq to determine the distribution of endometrial cell types across the secretory phase (n = 11), and in the decidua from first trimester pregnancies (n = 3) (**Table S1**). A total of 171,261 cells passed quality control with an average of 27,654 reads per cell. UMAP resolved 17 main cell clusters (**Figure 1D**). Cluster identities were assigned using the expression profiles of canonical markers for cell populations expected to be found in the non-pregnant human endometrium (**Figure 1E**). Cells were identified in all clusters across all groups, and the proportion of each cell type varied by stage (**Figure 1F and Figure S1B**). We identified similar cell clusters in the decidua, however, there were notable differences in the proportions of each cell type when compared to the secretory phases. Proliferative stromal cells were most abundant in the early secretory endometrium (89% of the cluster) whereas lymphatic endothelium (61%) and four clusters of immune cells, myeloid (50%), natural killer (NK) (50%), NK/Lymphoid (50%), and B cells (49%), were most abundant in the endometrial decidua of pregnancy. Additionally, compared to the decidua, all immune cell populations were decreased in the mid-secretory phase samples (early = 24%, mid = 7%, late = 22%, decidua = 42%). Taken together, the global transcriptome and the single-cell analysis demonstrated dynamic shifts in endometrial gene expression and cell composition across the menstrual cycle, with a marked transition from an early secretory to mid-secretory phase phenotype and an immune-cell dominated environment in early pregnancy. These findings highlight key molecular and cellular changes that may be critical for endometrial receptivity and successful embryo implantation.

### Expression of ESR1 and PGR in the epithelial and stromal compartments of the secretory phase endometrium and first trimester decidua

Ovarian-derived estradiol (E2) and progesterone (P4) bind to their cognate receptors, estrogen receptor alpha (ESR1) and progesterone receptor (PGR), which mediate hormonal signaling to prepare the uterine lining for embryo implantation by regulating cellular proliferation, differentiation, and receptivity^34,35^. Circulating E2 and P4 levels were measured at the time of endometrial sampling (**Table S1**), and we recently showed histopathological alignment of endometrial glandular and stromal compartments to menstrual cycle stage when the day of endometrial sampling was based on detection of the urinary LH surge^13^. To understand how circulating E2 and P4 could impact the endometrium, we defined the spatial and temporal changes in *ESR1* and *PGR* across stages of the menstrual cycle in the epithelium and stroma (**Figure 2**). *ESR1* expression was observed in both stromal and epithelial cells during the early secretory phase, with higher levels in the glandular and secretory glandular epithelium (**Figure 2A**). Compared to the early secretory phase, *ESR1* expression was reduced in the mid- and late secretory phases and became more restricted to the proliferative stroma and glandular epithelium. These findings were corroborated at the protein level (**Figure 2C**). Similarly, *PGR* was highly expressed in both stromal and epithelial cells during the early secretory phase (**Figure 2B**) but showed a marked decrease in the mid- and late secretory phases, with very low expression in the epithelium. This pattern was also supported at the protein level (**Figure 2D**). These data confirm the dynamic regulation of ESR1 and PGR expression, highlighting a shift from widespread hormonal responsiveness in the early secretory phase to a more localized and reduced expression pattern in the mid and late secretory phases. This transition likely reflects the endometrium’s preparation for implantation and its progression toward a refractory state. Moreover, comparison with the first trimester decidua revealed continued expression of PGR in the stroma and a further reduction in epithelial *ESR1* and *PGR* expression (**Figures 2C-D**).

**Figure 2.**
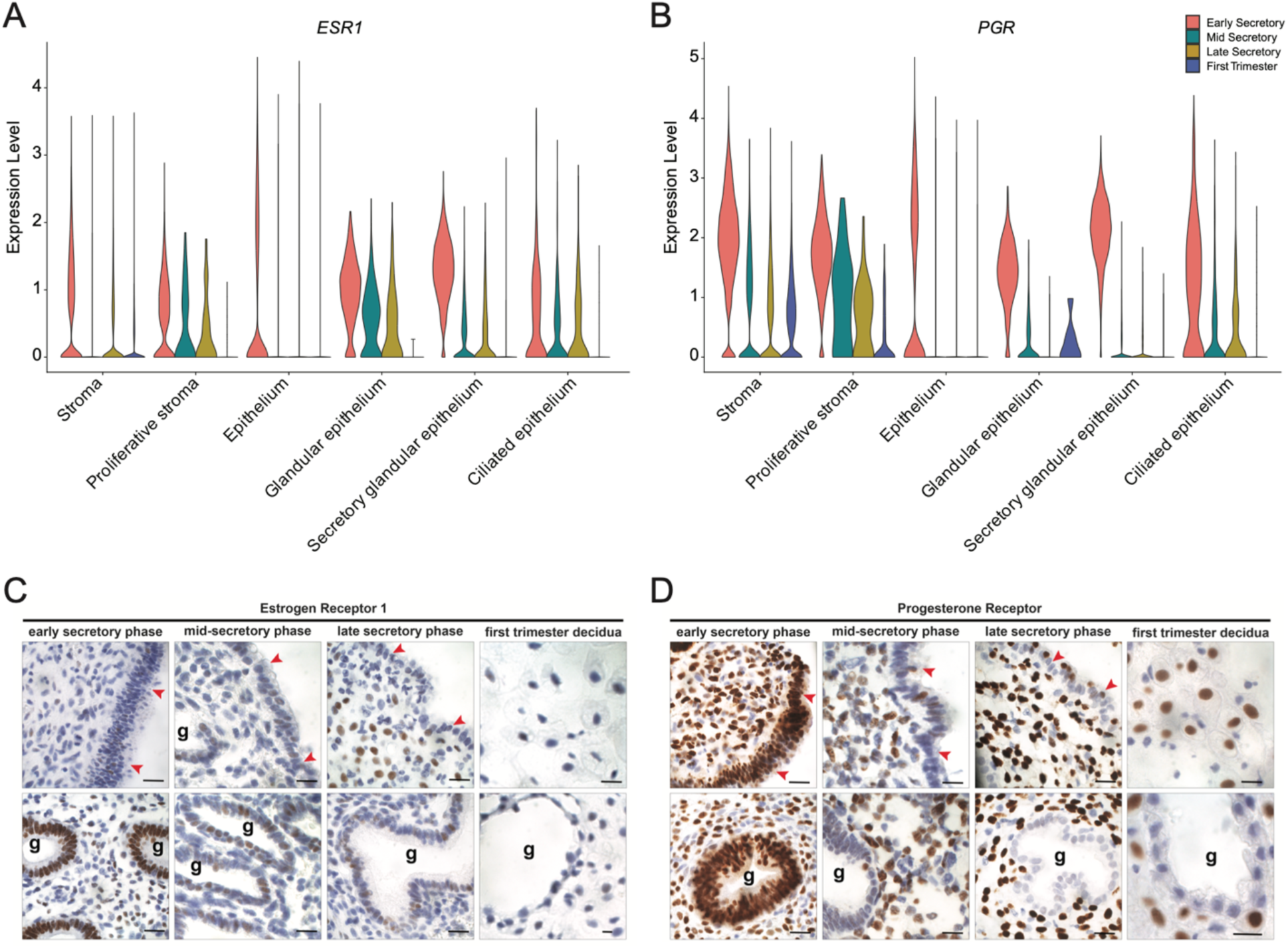
Expression of ESR1 and PGR in secretory phase endometrium and first trimester decidua. (A, B) Violin plots showing the expression of *ESR1* (A) and *PGR* (B) in the stromal and epithelial clusters across the secretory phase and in first trimester decidua. (C. D) Representative images of endometrial tissue sections stained to detect expression of ESR1 (C) and PGR (D) in glands and stroma of early, mid- and late secretory phase endometrium and endometrial decidua of first trimester pregnancy. g = gland; red arrowheads = luminal epithelium. Scale bars = 10 µm.

### Endometrial Receptivity Array (ERA) genes are expressed in the glandular epithelium

The expression pattern of 238 genes defines the Endometrial Receptivity Array (ERA), a diagnostic tool that was developed to identify the optimal timing for human embryo transfer based on an endometrial receptivity signature^20^. Analysis of gene expression patterns in our bulk mRNA-seq data, demonstrated that 84% of ERA genes were differentially expressed, in a consistent direction, from early compared to mid-secretory phase samples (132 increased, 68 decreased) and only 3% (4 increased, 3 decreased) overlapped with DEG from mid-compared to late secretory phase samples (**Figure 3A-C).** These findings confirmed expression of ERA genes in the mid secretory phase. Next, utilizing an “ERA score”, which was defined by the 143 genes that are upregulated in the ERA^20^, we used scRNA-seq analysis of the endometrial samples to identify cell types corresponding to the receptivity signature. Glandular epithelium and secretory glandular epithelium had the highest expression levels of ERA genes (**Figure 3D).** The ERA marker genes had the lowest expression in early secretory stage, with an overall increase at the mid-secretory phase, and a continued rise in expression in the secretory glandular epithelium of late secretory phase and first trimester decidua samples (**Figure 3E**). Taken together, these data suggest that while the ERA gene signature is enriched in the mid-secretory phase, its expression extends beyond the traditional implantation window, persisting into the late secretory phase and early pregnancy. This finding highlights the power of scRNA-seq for understanding temporal changes in gene expression within distinct epithelial cell types.

**Figure 3.**
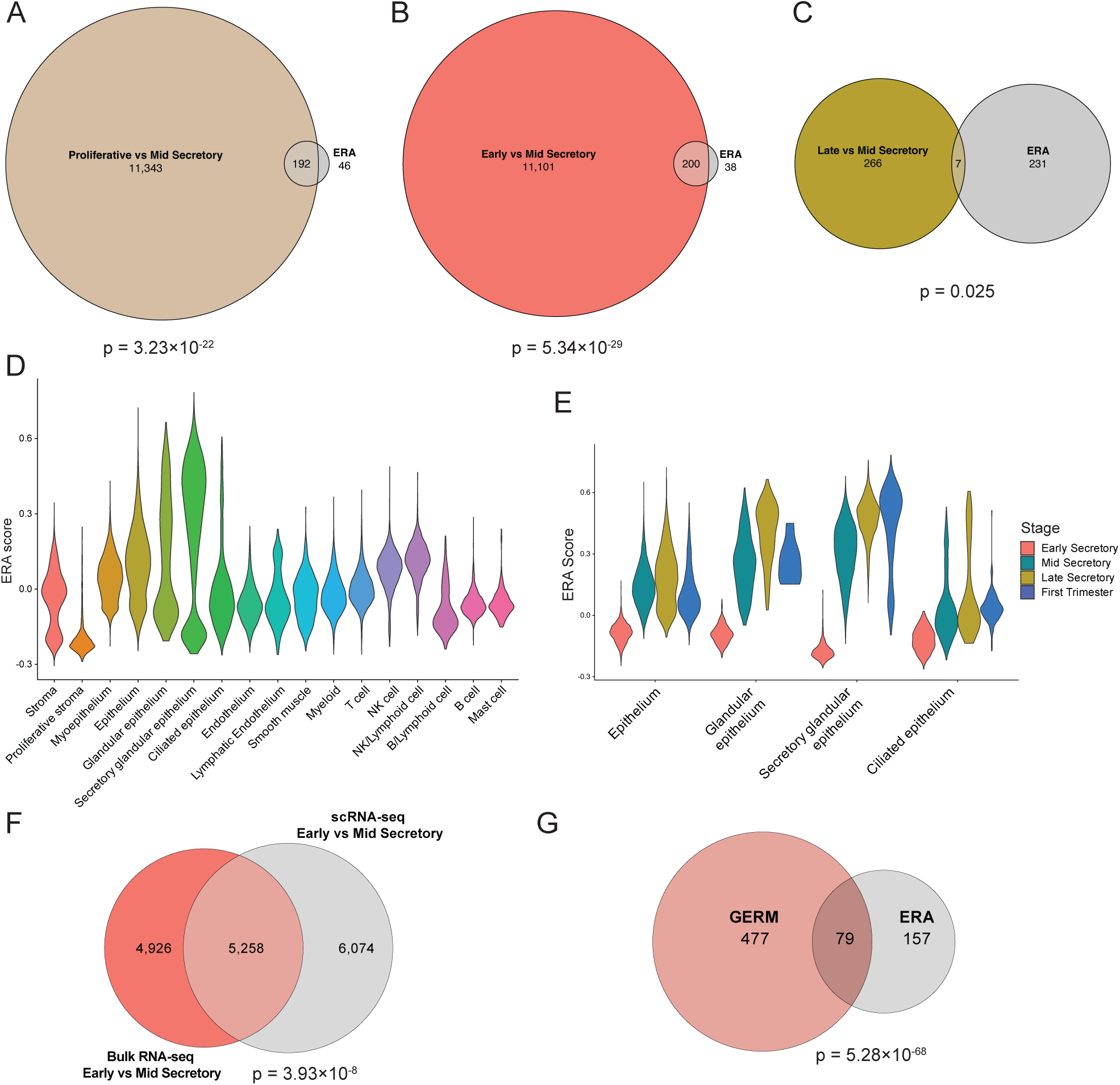
Endometrial receptivity array (ERA) gene expression in bulk mRNA-seq and scRNA-seq analyses. (A-C) Euler plots of genes with significantly different expression in the proliferative vs mid-secretory (A), early vs mid-secretory (B), and late vs mid-secretory (C) phases as compared to the ERA. Hypergeometric test p-values are indicated for each comparison. (D, E) Violin plots of the ERA score in each cell cluster (D) and the epithelium clusters across stages (E). (F) Overlap of DEG from bulk and scRNA-seq secretory glandular epithelium comparing early to mid-secretory endometrium. (G) The glandular epithelium receptivity module (GERM) score genes include 79 (33%) of the ERA genes.

Based on the dispersion of the epithelial cluster in the UMAP analysis, we performed a subcluster analysis. In addition to glandular, secretory glandular, and ciliated epithelium, we identified seven epithelial subclusters (**Figure S2A**). The proportion of cells within each cluster varied dynamically, with an expansion of cells in epithelium 0 from the early secretory phase to later stages and a decline of cells in glandular and secretory glandular epithelium in first trimester decidua samples (**Figure S2B**). These findings reveal stage-specific epithelial remodeling, potentially driven by shifts in cellular composition or dynamic transcriptional states.

To identify marker genes that could define a receptive transcriptomic signature of the mid-secretory endometrium, we identified DEG in the secretory glandular epithelial cells in early versus mid-secretory samples. 11,332 genes were differentially expressed (**Table S6**) with 5,258 overlapping DEG identified in bulk RNA-seq comparisons of early vs mid-secretory (**Figure 3F**). We further filtered these overlapping DEG using a cutoff of log_2_ fold-change > |2| and an FDR < 10^-7^ and found 556 marker genes (267 down- and 287 up-regulated, **Table S7**) that we termed the glandular epithelium receptivity module (GERM). Notably, 33% of the ERA (n = 79) genes were included in the GERM signature (**Figure 3G**).

### *In vitro* decidualization and senescent marker genes are highly expressed in decidua from first trimester pregnancy

*In vivo,* estrogen primed endometrial cells differentiate under the influence of progesterone, thus generating an endometrium that is receptive to embryo implantation. Stromal cell decidualization, which is an integral part of endometrial remodeling, has been frequently studied, perturbed or recovered *in vitro* to gain a deeper understanding of the mechanisms underlying appropriate versus aberrant endometrial differentiation. Decidualization of stromal cells *in vitro* is commonly verified by increased expression of “classic” markers of decidualization including *IGFBP1*^36^*, PRL*^37^ *and FOXO1*^38,39^. We combined these “classic” markers, as well as others associated with *in vitro* decidualization^40,41^ and found that expression of genes associated with *in vitro* decidualization was highest in the first trimester decidua when visualized across the secretory phase and first trimester samples (**Figure 4A**). The gene list was then used to generate a “decidualization score” to identify cells that expressed *in vitro* decidualization marker genes. Cells exhibiting high marker expression were predominantly found in the first trimester stromal cluster, with lowest expression observed during the early secretory phase compared to the mid and late secretory phases (**Figure 4B**). Stromal cells with a rounded, epithelioid morphology and prominent nuclei, as is seen after *in vitro* decidualization of isolated primary or immortalized endometrial stromal cells^42^, were detected by H&E staining in the first trimester, but not in the mid-secretory phase (**Figure 4C**). These data demonstrate gene expression and the differentiated cellular morphology consistent with decidualization in stromal cells in first trimester deciduae.

**Figure 4:**
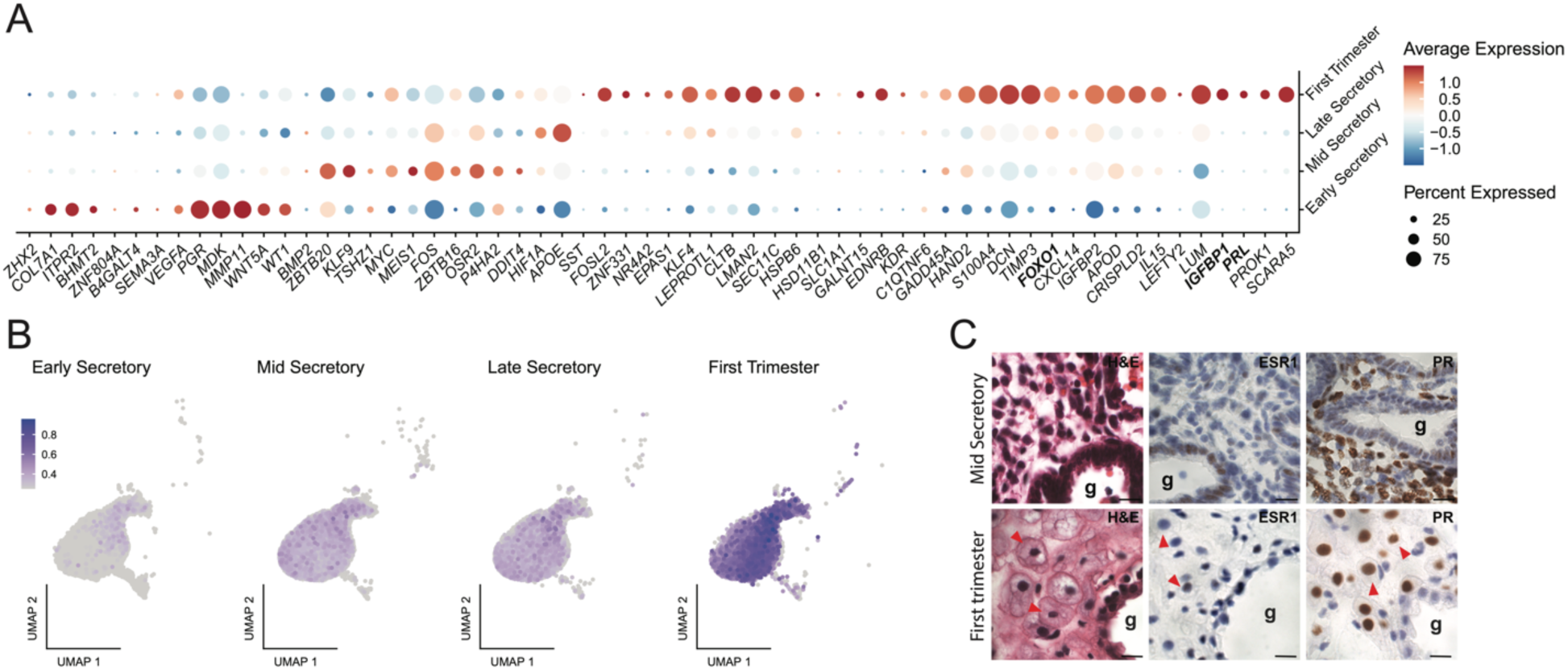
Markers of *in vitro* decidualization are highly expressed in endometrial decidua from the first trimester of pregnancy. (A) Dotplot of *in vitro* decidualization markers in the stroma cluster separated by stage. Expression of *IGFBP1, FOXO1* and *PRL* is highlighted with bold type. (B) UMAP of the *in vitro* decidualization marker score in the stroma cluster. Average score expression is represented by color intensity. (C) Representative images of tissue sections from the mid secretory phase and first trimester decidua stained with H&E g = gland; red arrowheads = decidualized stromal cells with epithelioid morphology. Scale bars for H&E = 10 µm.

Cells with an elevated decidualization score were not uniformly distributed within the stromal cluster, supporting the utility of re-clustering the stromal population for more detailed analyses. Six stromal subclusters were found in addition to the previously labeled proliferative stroma cluster (**Figure 5A**). The cell type proportions varied across stages, except stroma 4 (early = 11%, mid = 15%, late = 11%), with an increase from the early to mid-secretory and late secretory phases in stroma 1, 3 and 5 (Stroma 1: early = 11%, mid = 24%, late = 29%; Stroma 3: early = 13%, mid = 23%, late = 21%; Stroma 5: early = 0.1%, mid = 0.6%, late = 0.3%). This corresponded to decreases in proliferative stroma and stroma 0 and 2 (Proliferative: early = 8%, mid = 0.3%, late = 0.3%; stroma 0: early = 33%, mid = 20%, late = 24%; stroma 2: early = 24%, mid = 18%, late = 14%) (**Figure 5B-C**). Analysis of *in vitro* decidualization markers in the subclusters confirmed higher expression in the first trimester stromal cells (**Figure 5D**). However, no specific cell population demonstrated consistently elevated expression of these *in vitro* markers (**Figure 5D-E**).

**Figure 5:**
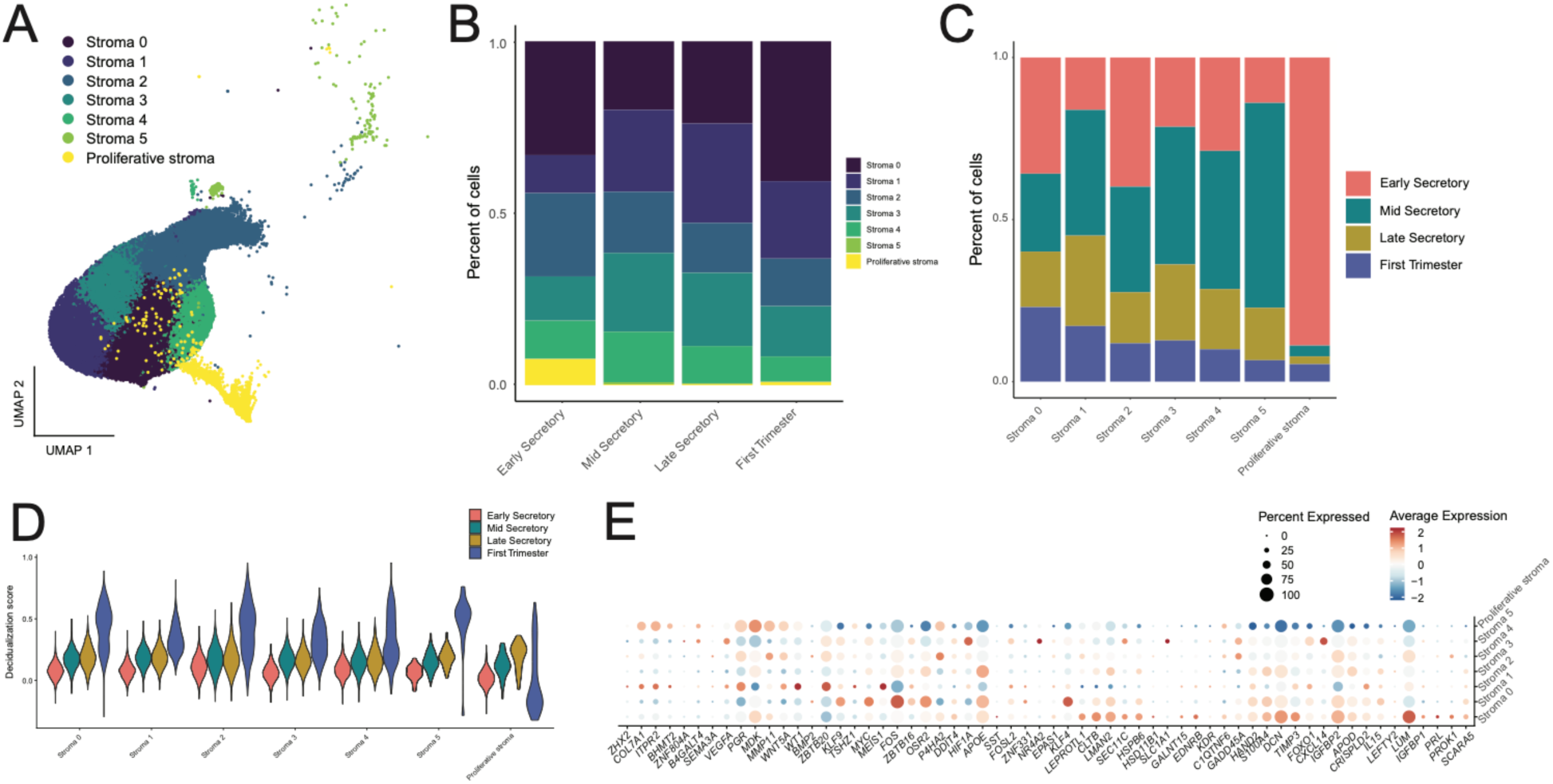
Stromal subcluster distribution and diffuse expression of *in vitro* decidualization markers in secretory phase endometrium and first trimester decidua. (A) UMAP of stroma subclustering showing five stroma subclusters in addition to the proliferative cluster. (B) Stacked bar plot of the stroma cells showing percentages of cells in the stroma subclusters by stage. (C) Stacked bar plot of the stroma cells showing percentages of cells for each stage by subcluster. (D) Violin plot of the decidualization score across the stroma subclusters split by stage. (E) Dotplot of *in vitro* decidualization markers in the stroma subclusters.

### Senescent stroma cells are the most abundant in the secretory phase endometrium and decidua from first trimester pregnancy

Senescent stromal cells are present in secretory phase endometrium and likely play important roles in fertility^40^. We utilized a panel of *in vitro* cellular senescence marker genes^40,41^ to identify senescent cells in our endometrial samples. We found 7 of the 12 genes had the highest expression in first trimester endometrial deciduae (Figure 6A), mirroring expression of in vitro decidualization marker genes, and that expression of *in vitro* senescence markers was not restricted to a single stroma subcluster (**Figure S3**). We then used expression of validated *in vivo* markers, *SCARA5* and *DIO2*^40^, to differentiate decidual and senescent subclusters. Stroma subcluster 1, 3 and 5, which are most abundant in the mid- and late secretory phase samples, had highest expression of *SCARA5*, and were identified as decidualized stromal cells (DSC) (**Figure 6B**). These were further characterized by expression of *FOS*, denoted as DSC *FOS*^lo^ and DSC *FOS*^hi^, or high expression of *CXCL14*, denoted as DSC *CXCL14*^hi^ (**Figure 5E and 6B**). Stroma 0 and 4, with increased expression of *DIO2*, were identified as senescent stroma subclusters and were further characterized by expression of *DIO2*, denoted as *DIO2*^lo^ and *DIO2*^hi^ (**Figure 6B**). Stroma subcluster 2 expressed *SCARA5* and *DIO2* and was labeled senescent decidualized stromal cells (snDSC) (**Figure 6B**).

**Figure 6:**
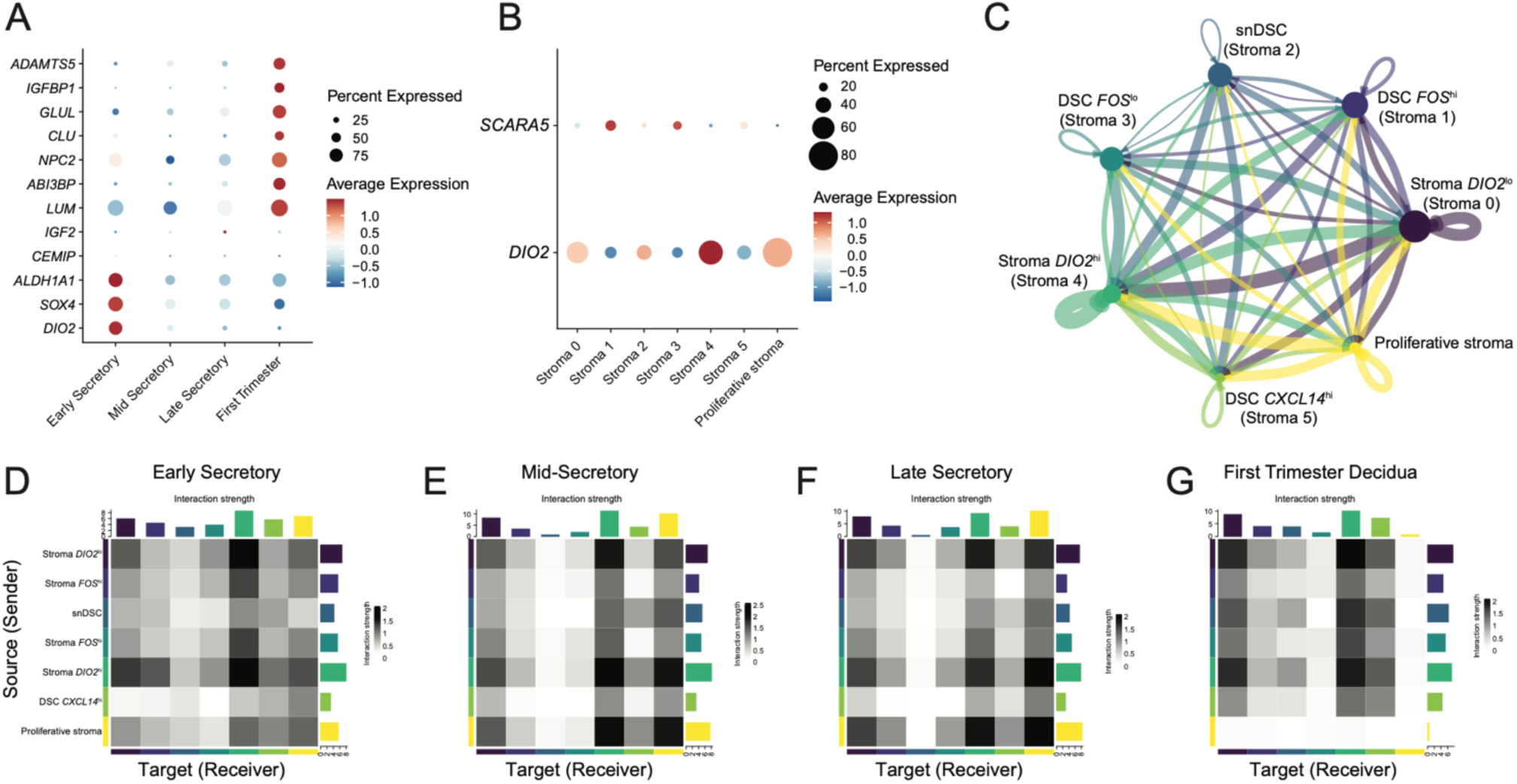
Cluster identification and communication in the stroma. (A) Dotplot of *in vitro* senescence markers across stages. (B) Dotplot of *SCARA5* and *DIO2*, *in vivo* markers for decidualization and senescence, respectively. (C) Cell-cell communication of the stroma clusters. Edge colors are consistent with the sources as sender, and edge weights are proportional to the interaction strength, that is a thicker line indicates a stronger signal. Circle sizes are proportional to the number of cells in each cluster. Communication in the early secretory (D), mid-secretory (E), and late secretory (F) phases, and first trimester decidua of pregnancy (G) are shown as heatmaps. Colored bars represent the relative signaling strength of pathway across subclusters. The top-colored bar plot represents the sum of each column of the absolute values displayed in the heatmap (incoming signaling). The right colored bar plot represents the sum of each row of the absolute values (outgoing signaling).

To identify unique signaling patterns among stromal subclusters, ligand-receptor communication analysis was conducted using CellChat^43,44^. This analysis revealed that overall stromal communication was driven by senescent *DIO2*^lo^ and *DIO2*^hi^ cells (**Figure 6C**). In contrast, the snDSC cluster displayed lowest overall signaling (**Figure 6C**). Assessment of stromal cell-cell communication across stages showed similar signaling patterns in mid- and late secretory phases with snDSCs having the lowest receiver score (**Figure 6E-F**). Additionally, increased communication between the proliferative stroma and senescent *DIO2*^hi^ cells was evident during mid- and late secretory phases (**Figure 6E-F**). Based on sender and receiver scores, senescent *DIO2*^hi^ and *DIO2*^lo^ and proliferative stroma cells were the most interactive clusters, during early, mid-, and late secretory phases (**Figure 6D-F**). In contrast, proliferative stroma was the least interactive subcluster in first trimester deciduae (**Figure 6G**).

### Stromal cell communication with secretory glandular epithelium increases during the mid-secretory phase

The epithelial and stromal compartments of the endometrium have independent and interdependent roles in supporting embryo implantation^45,46^. While the epithelium undergoes critical modifications to establish receptivity for nidation^47,48^, stromal cell differentiation is essential for successful implantation and subsequent pregnancy maintenance^49^. However, the mechanisms by which epithelial and stromal cells coordinate their functions during this process remain incompletely defined. To better understand the interactions between these two compartments, we defined the crosstalk between the epithelium and stroma cell populations.

Epithelial clusters 0 through 4 had the lowest receiver strength across all stages and had low interaction strengths. In the secretory phases, the glandular, secretory glandular and all stroma sub-clusters were interactive (**Figure 7A-C**). Epithelial cells exhibited reduced cell-cell communication in the first trimester decidua samples. In contrast, signaling between all stroma subclusters except for the proliferative stroma was active in the first trimester decidua samples (**Figure 7C**).

**Figure 7:**
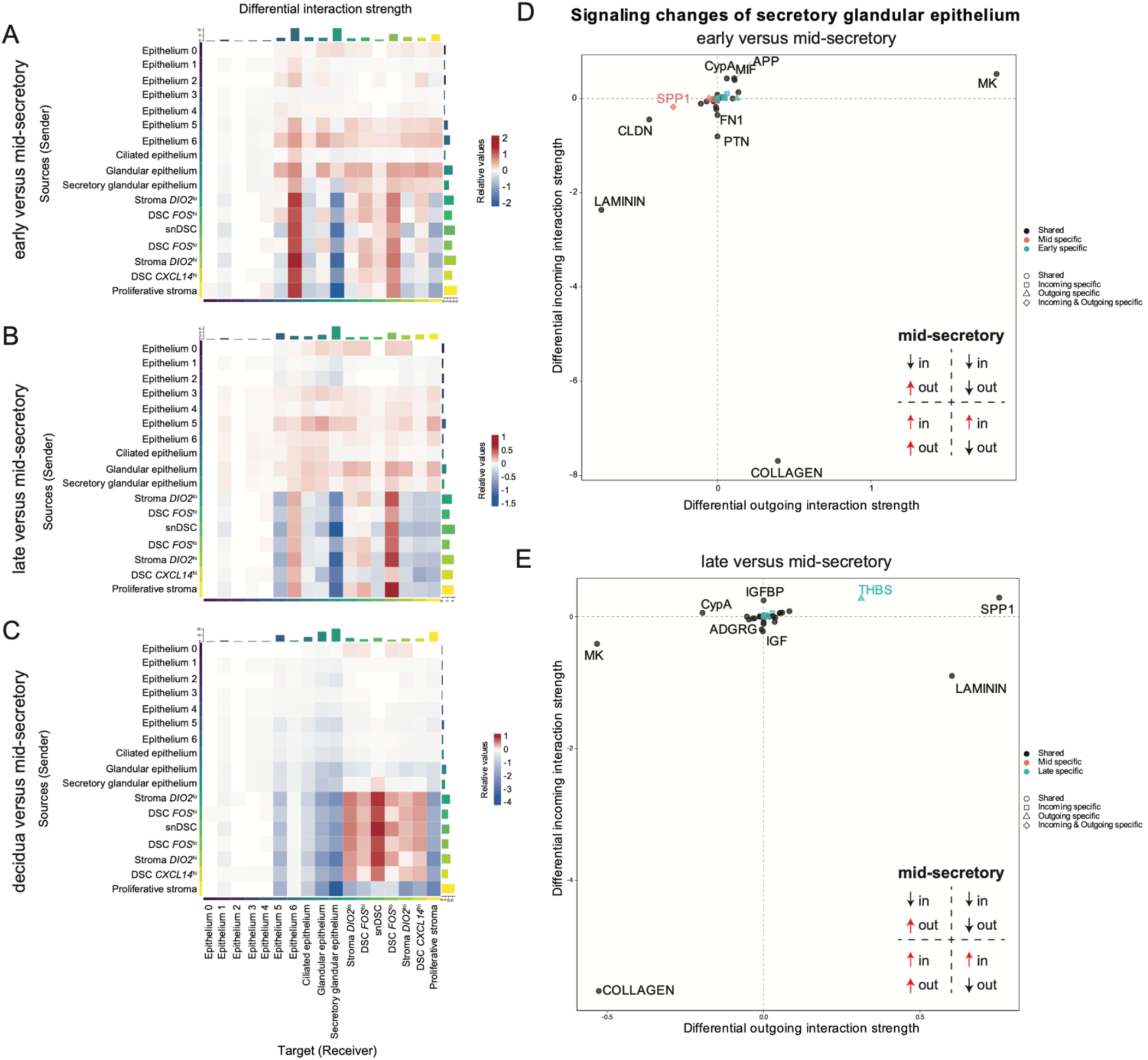
Stroma-epithelium communication peaks at mid-secretory phase in the secretory glandular epithelium. (A-C) Communication changes between the epithelium and stroma clusters in the early vs mid-secretory phase (A), late vs mid-secretory phase (B), and first trimester decidua vs mid-secretory phase (C) shown using heatmaps. Colors represent the relative signaling strength of a signaling pathway across clusters with the blue indicating decreased communication and the red an increase of communication probability. The top-colored bar plot represents the sum of the absolute values for each column displayed in the heatmap (incoming signaling). The right colored bar plot represents the sum of the absolute values of each row (outgoing signaling). (D, E) Signaling changes in the secretory glandular epithelium cluster is shown in scatter plots for early versus mid-secretory (D) and late versus mid-secretory (E).

Epithelium 6 and DSC *FOS*^lo^ clusters had higher receiver strength in the early secretory compared to the mid-secretory phase, with stroma as the predominant source. By contrast, the stroma clusters had reduced communication with the secretory glandular epithelium in the early secretory phase samples (**Figure 7A**). A similar pattern was observed for late secretory compared to mid-secretory phase samples (**Figure 7B**), indicating that stroma to secretory glandular epithelium communication was increased during the mid-secretory phase. Conversely, communication from the stroma to epithelium 6 and DSC *FOS*^lo^ was decreased in the mid-secretory phase samples (**Figure 7A-B**). In the first trimester deciduae compared to mid-secretory phase samples, all stroma subclusters, except proliferative, had increased interaction strength within the stroma (**Figure 7C**). Interestingly, receiver strength for the snDSC cluster was increased in the first trimester decidua samples compared to mid-secretory phase samples.

Since the transition from the early to mid-secretory phase is critical for pregnancy initiation, we investigated altered signaling pathways in the secretory glandular epithelium, the primary cell type exhibiting changes in communication between the epithelium and stroma. Pathways associated with the extracellular matrix, including collagen and laminin, were downregulated in the secretory glandular epithelium of early and late secretory phase samples compared to mid-secretory phase samples (**Figure 7D-E**). Signaling network analysis during the mid-secretory phase confirmed that the secretory glandular epithelium was the strongest signal recipient within the collagen pathway, receiving input from all stromal subclusters, as well as the epithelium 5 and ciliated epithelium subclusters (**Figure 8A**). The ligand-receptor interaction was strongest during the mid-secretory phase (**Figure 8C**) compared to the early (**Figure 8B**) and late secretory phases (**Figure 8D**), suggesting a key role for collagen signaling in endometrial receptivity. Further analysis of ligand-receptor interactions within the collagen pathway revealed that the primary ligand contributors were *COL1A1*, *COL1A2*, *COL6A1*, *COL6A2*, *COL4A1* and *COL4A2* while *CD44* was the predominant receptor in the secretory glandular epithelium (**Figure 8E**).

**Figure 8:**
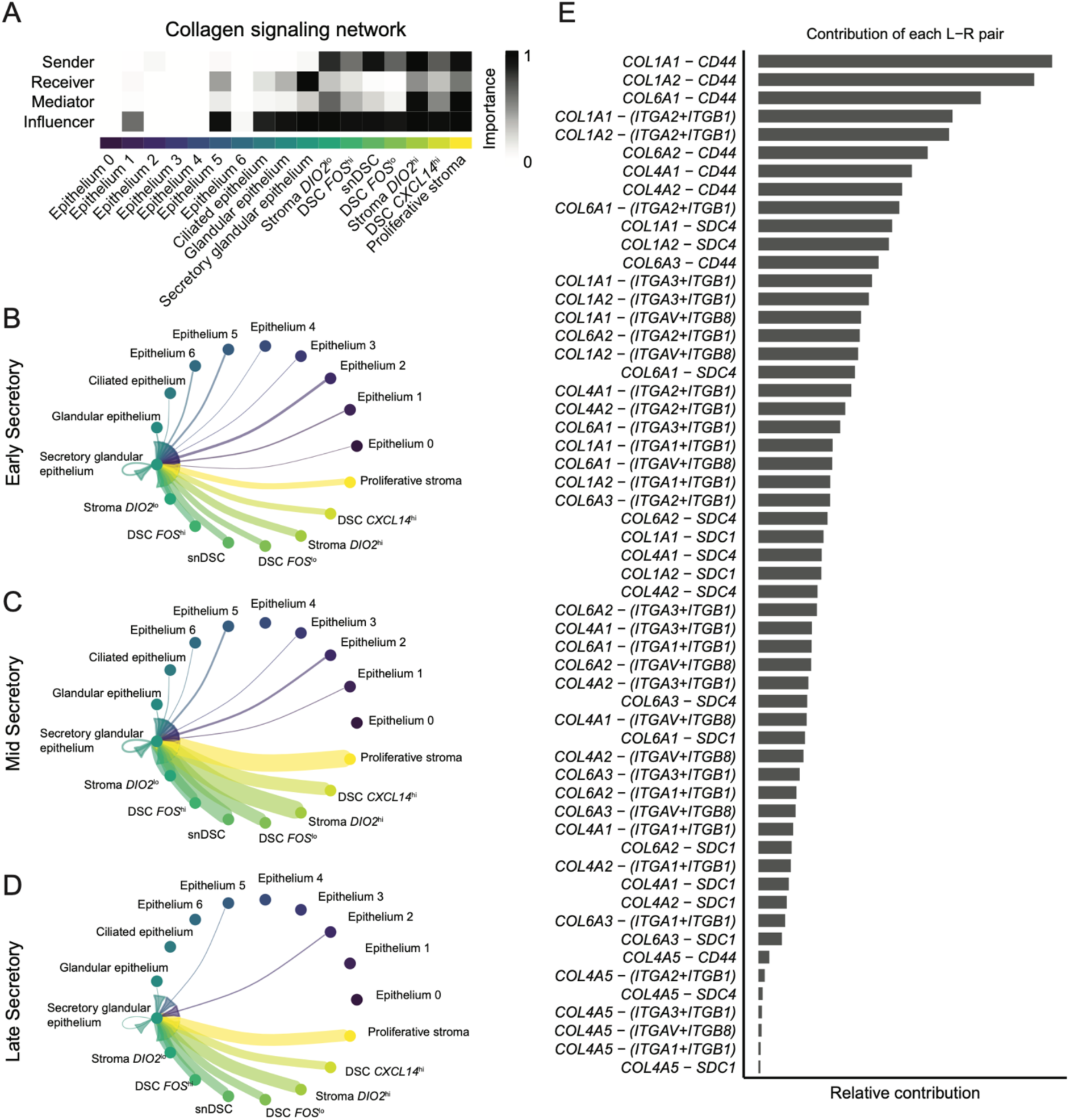
Collagen signaling between the stroma and epithelium during the secretory phase. (A) Role of each cell cluster in collagen signaling shown as a heatmap. (B-D) Cell-cell communication of the stroma and epithelium with the secretory glandular epithelium as the recipient. Edge colors are consistent with the sources as sender, and edge weights are proportional to the interaction strength, that is a thicker line indicates a stronger signal. Circle sizes are proportional to the number of cells in each cluster for the early secretory (B), mid-secretory (C), and late secretory (D) phases. Note the increased communication from stroma subclusters during mid-secretory and late secretory phases. (E) Relative contribution of ligand-receptor pairs for signals coming into the secretory glandular epithelium.

### Dysregulated expression of epithelial- and stromal-cell specific genes in the endometrium of patients with infertility

Based on the observed temporal increase in communication between stroma and secretory glandular epithelium during the mid-secretory phase, as well as the localization of genes associated with endometrial receptivity within the same epithelium, we hypothesized that this signaling axis would be altered in endometrium of patients suffering from infertility. We first identified decidual and senescent stromal cells in a publicly available dataset possessing endometrial samples at the mid-secretory stage from controls and patients with recurrent implantation failure (RIF) (GSE183837)^2^. Stromal populations were classified in control samples based on our scRNA-seq markers (**Figure 9A**) and we found disrupted proportions in endometrial samples from RIF patients (**Figure 9B**). There was an expansion of senescent cells, expressing *DIO2*, and a concurrent decrease in decidual cell populations. Notably, *CXCL14*^hi^ decidual stromal cells were severely decreased in samples from RIF patients (0.2% vs 30%).

**Figure 9.**
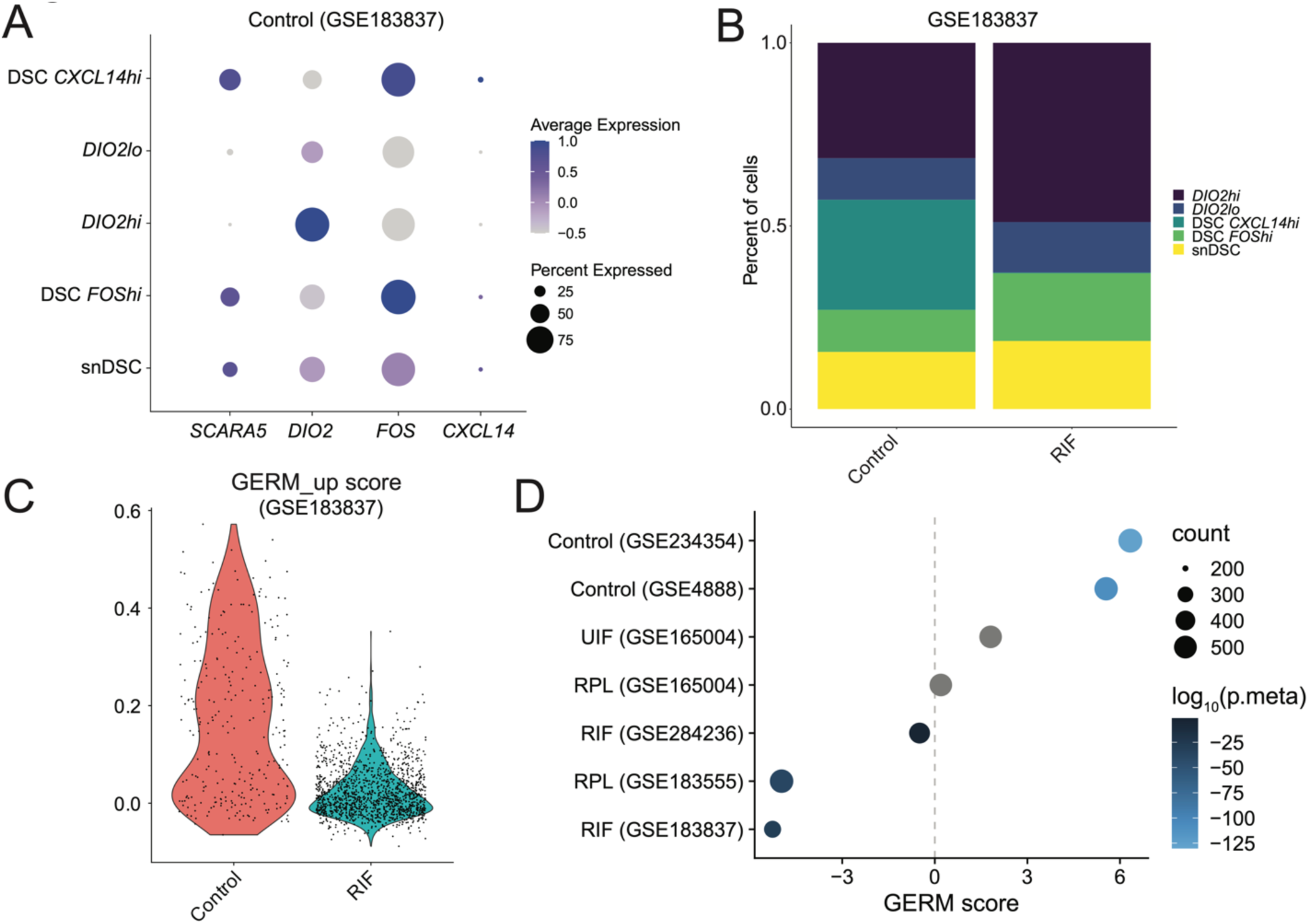
Disrupted stromal cell populations and epithelial gene expression in patients with infertility. (A) Dotplot of *SCARA5*, *DIO2*, *FOS*, and *CXCL14* for identification of the stroma populations in control endometrium samples from a scRNA-seq dataset (GSE183837^2^) (B)Stacked bar plot shows altered stroma cell proportions in endometrial samples from RIF patients. (C) Glandular epithelium receptivity module (GERM) upregulated genes in the secretory glandular epithelium are reduced in RIF patients. (D) GERM score is positively correlated with mid-secretory endometrium from control patients (GSE234354^33^ and GSE4888^50^), but not from patients with unexplained infertility (UIF, GSE165004^52^), recurrent pregnancy loss (RPL, GSE165004 and GSE183555^53^), or RIF (GSE284236^51^ and GSE183837^2^) A gray circle indicates an insignificant combined p-value, or p.meta ≥ 0.05.

We then applied our GERM signature to multiple datasets including those containing endometrial early and mid-secretory phase samples from fertile controls^33,50^ and patients with recurrent implantation failure^2,51^ (RIF), recurrent pregnancy loss^52,53^ (RPL), and unexplained implantation failure^52^ (UIF) **(Figure 9C-D**). We confirmed reduced expression of GERM signature genes (GERM_up only) in the secretory glandular epithelium in endometrial samples from RIF patients (**Figure 9C**). Gene set enrichment analyses (GSEA) using our GERM signature, GERM_up and GERM_down, were then combined to produce a GERM score for the other datasets, which included microarray, bulk RNA-seq, scRNA-seq, and spatial RNA-seq results. The GERM score was significant and positively correlated (scores = 5.6-6.3) with healthy endometrium in the mid-versus early secretory phase **(Figure 9D**). Our bulk RNA-seq dataset, for comparison, had a GERM score of 8 with a combined p-value of 0 as a positive control. In contrast, GERM scores in endometrial samples of patients with infertility, specifically RIF, RPL, or UIF, were decreased (scores = -5.3-1.8). The GERM score from one microarray study^52^ (GSE165004) was reduced (scores = 0.2-1.8), but not significantly different. Inspection of the fold changes revealed a limited dynamic range that likely affected the strength of the GSEA analysis. Together, these findings validate of our GERM score across control patients of different racial and demographic origins and demonstrates dysregulation our GERM genes in patients with “endometrial factor” infertility, The GERM gene list expands the panel of genes associated with endometrial receptivity, specifically capturing those expressed in secretory glandular epithelial cells, and supports our finding that the glandular epithelium gene signature is correlated with successful embryo implantation

## Discussion

In this study, we performed bulk and single-cell transcriptomic analyses on endometrial samples from ovulatory menstrual cycles and deciduae from first-trimester pregnancies to elucidate the epithelial and stromal changes driving endometrial differentiation in preparation for implantation. This is the first study to include fertile Black and Hispanic patients, addressing a critical gap in reproductive research. We show that glandular epithelial cells are central to endometrial receptivity and shifts in stromal decidual and senescent populations, highlighting the complex, multi-lineage endometrial changes that underlie embryo implantation.

Estrogen receptors and progesterone receptors mediate hormone signaling pathways that are essential for regulating changes in the endometrium throughout the menstrual cycle in preparation for potential pregnancy. Comparison with prior studies demonstrated that ESR1 and PGR gene and protein expression^54,55^, as well as gene expression profiles in the epithelial compartment^33,50^, are consistent among fertile and presumably fertile patients with regular menstrual cycles. The distribution of ESR1 and PGR proteins in our secretory phase samples aligns with published literature, which indicates that ESR1 and PGR expression is highest in the proliferative phase and decreases as the menstrual cycle progresses^56–60^. This finding, along with finding similar gene expression in control patients of different racial and demographic origins suggests that the molecular signature in the window of implantation is shared across racial and ethnic groups - an important foundation to establish when comparing our data with previously published data on patients with infertility.

Since at least the early 20^th^ century, clinicians and scientists have been attempting to describe and characterize the features of an endometrium that is receptive to embryo implantation. In 1950 Noyes et al put forth criteria for dating endometrial biopsies based on phenotypic features of endometrial glands and stroma on histopathology^37^, facilitating the uniform categorization of a biopsy sample as early-, mid- or late-proliferative, and describing the secretory phase in terms of number days post-ovulation. Noyes criteria was later applied to the examination of luteal phase defects^40^ and assessed for its correlation with fertility status^41^ but has been largely relegated to the realm of pathologists for descriptive purposes only. More recently, attempts were made to characterize the window of implantation based on a transcriptomic signature, a technique theorized to optimize embryo transfer timing by uncovering pre-receptive and post-receptive endometria on the day of typical receptivity^17^. This 238-gene tool, known as the endometrial receptivity array (ERA), did not demonstrate an improvement in pregnancy or live birth rates^18^. Despite the limitations of the ERA in improving pregnancy rates, we were interested in examining the markers of receptivity used in their analyses to understand the cellular origins of the receptivity signature and to explore the cellular and molecular factors that contribute endometrial receptivity. In this study, we found that the glandular epithelium cluster had the greatest expression of ERA genes, emphasizing the importance of this cellular compartment at the embryo-endometrium interface. Previous work has suggested that the molecular signature of the glandular epithelium correlates strongly with menstrual cycle day^61^. Our results expand on this finding by suggesting that the importance of the glandular epithelium lies in its key role as the cell type driving receptivity in the mid-secretory endometrium. By identifying glandular epithelial markers of receptivity that are detectable in bulk RNA sequencing we generated a set of receptivity markers (the *Glandular Epithelium Receptivity Module*, or GERM score) which was applied to published datasets and found to be altered in conditions suggestive of abnormal endometrial differentiation such as RIF and RPL. This novel score therefore represents a promising new direction in the characterization of endometrial receptivity.

Based on our data, we have proposed an important role for the glandular epithelium in defining endometrial differentiation and generating a receptivity signal. Understanding that endometrial differentiation is highly dependent on stromal cell decidualization as well, we next sought to examine this cellular compartment in greater depth. We compared endometrial gene expression to the gene expression patterns of *in vitro* stromal cells and uncovered key differences between these two models. Prior studies have examined the characteristics of stromal cells *in vitro* to gain a better understanding of the properties of these cells before and after the addition of a decidualization stimulus. *In vitro* decidualization is commonly carried out by culturing human endometrial stromal cells (HESCs) with one or more of the following stimuli: cyclic AMP (cAMP), medroxyprogesterone acetate (MPA) and estradiol^62^. In these studies, decidualization is typically confirmed by appreciating changes in cellular morphology, from fibroblast-like to epithelioid, and assessing expression of markers including insulin-like growth factor binding protein 1 (IGFBP1) and prolactin (PRL)^62^. When we examined the expression of IGFBP1 and PRL in our secretory phase and first trimester decidua samples we found the highest expression of these markers in the pregnancy decidua, with relatively lower levels of expression in the secretory phase, suggesting that these markers may be used more aptly in models of early pregnancy decidua rather than to mimic the secretory phase endometrium in the absence of embryo implantation. To our knowledge, our study is one of the first to directly compare cell-type specific expression of these two markers and cellular morphology in both secretory phase biopsies and pregnancy decidua. Our data are in line with those that suggest that in the absence of pregnancy the secretory phase can be thought of more aptly as a “pre-decidua” that only achieves full decidualization after embryo implantation^63^. Our findings suggest the need for more complex *in vitro* assays to better assess the interactions between cells, as well as the need for a stimulus that more closely mimics the differentiation of the secretory-phase endometrium. In this way, the pre-implantation endometrium may be studied ex vivo with higher fidelity.

Cellular senescence refers to the permanent proliferative arrest of a cell in response to various stressors^64^. This process has been described in two forms: acute (which is transient and physiological) and chronic (persistent and age-related)^65,66^. Acute senescence may be observed as a part of normal processes, including embryogenesis, endometrial cycling, and repair of tissue injury^67,68^, whereas chronic senescence may represent the age-related or pathologic decline in tissue function^67,68^.There is a growing appreciation for the role of senescence in endometrial remodeling^66,69–73^. Experimental evidence indicates that HESC decidualization is accompanied by the appearance of a p16-positive senescent cell subpopulation, suggesting that cellular senescence is a critical component of normal HESC decidualization^71^. Embryo implantation may also activate physiologic senescence^74^. During decidualization, endometrial stromal cells undergo proliferation arrest and secrete inflammatory mediators, including senescence-associated secretory phenotype (SASP)^73^. While decidual senescence is critical for the initial pro-inflammatory response required for embryo implantation^75^, it is thought that premature senescence of human endometrial stromal cells can impair decidualization^70^. Within our six distinct stromal clusters, we resolved three clusters characterized by relative expression of decidualization marker *SCARA5* and senescence marker *DIO2*. Prior work has demonstrated an increase in senescent cells in the mid-secretory compared to the early secretory phase^61^, as well as increased expression of *DIO2* and decreased expression of *SCARA5* in subjects with recurrent pregnancy loss, compared to controls^40^. We found that the highest levels of stroma-stroma communication occurred in the senescent clusters, followed by the decidualized clusters, and the lowest levels of signaling occurred in the senescent-decidualized cluster. We therefore hypothesized that the relative proportions of cells in each subgroup may vary in normal and pathologic endometria and examined the expression patterns of *SCARA5* and *DIO2* in a published dataset that included women with RIF and controls. We found diminished expression of *DIO2* in women with RIF, as well as a larger proportion of cells in the senescent-decidualized cluster, suggesting that dysregulated senescence may contribute to, or be reflective of, suboptimal endometrial receptivity.

Our results highlight the dynamic crosstalk between stromal cells and the secretory glandular epithelium, particularly during the mid-secretory phase, underscoring the importance of these two cell types for endometrial receptivity. We observed that stromal-to-epithelial communication was highest in the mid-secretory phase, with the secretory glandular epithelium being the primary recipient of collagen-associated signaling. The extracellular matrix (ECM), particularly via collagen signaling, plays a fundamental role in embryo implantation by regulating endometrial tissue stiffness and mechanosensing through receptors such as CD44^76^. Interestingly, CD44 expression is reduced in infertile patients^77^, though mouse models lacking CD44 remain viable and fertile^78^, suggesting the presence of redundant receptors compensating for ECM-mediated signaling. This redundancy could be critical for maintaining implantation competence despite variations in individual receptor expression. The role of ECM mechanics in implantation is further supported by findings that collagenase-mediated softening of the ECM enhances fertility in mice^79^, reinforcing the hypothesis that ECM remodeling influences implantation efficiency.

The importance of epithelial glands in implantation has been well-documented across multiple species, including mouse^29–31^ and sheep^27,28^. Deletion of *Foxa2*, a glandular epithelium marker, demonstrated that the absence of this transcription factor lead to defective uterine gland development^80^ or function and subsequent implantation failure, underscoring the critical role of glandular secretions in establishing a receptive endometrial environment^81,82^. More recently ESR1-dependent uterine gland structure has been shown to be critical for production of the key glandular secretion Leukemia Inhibitory Factor^83^. Further, in the mouse, uterine glands show a characteristic reorganization of the branched glands towards the implantation site prior to embryo implantation supporting a dynamic role of uterine glands in uterine receptivity^84^. Similarly, studies in sheep indicate that endometrial glands produce key factors required for early pregnancy establishment, with disruption of glandular function leading to implantation failure^85^. These findings emphasize the conserved role of endometrial epithelial glands in implantation across species.

A limitation of our study is that it does not account for immune cell interactions, which are known to play a critical role in endometrial remodeling in preparation for embryo implantation. The maternal immune system modulates endometrial receptivity, and immune cells such as decidual macrophages and uterine natural killer cells are involved in ECM remodeling and trophoblast invasion. Recent studies highlighted the importance of immune-endometrial crosstalk in endometrium showing a dysregulating signaling in endometriosis particularly epithelium to macrophage through cytokine-mediated signaling pathways^51,86^. Future studies should integrate immune cell populations into cell-cell interaction models to provide a more comprehensive understanding of implantation dynamics.

## Methods

### Human Subjects

The use of human tissue specimens was approved by the Institutional Review Board at Rutgers Health (Pro2018002041). Self-identified Black and Hispanic women, aged 23-41 years, with regular, ovulatory menstrual cycles and at least one prior pregnancy were prospectively enrolled between November 2020 and December 2022. All study participants provided signed informed consent for the review of their medical records and collection of blood and endometrial samples. Women with endocrine or autoimmune disorders, including antiphospholipid syndrome and polycystic ovary syndrome, or anatomic disorders of the reproductive tract, including hydrosalpinges or submucosal fibroids, were excluded. Women with a history of pregnancy complications, including recurrent pregnancy loss, preeclampsia or intrauterine growth restriction were also excluded. The demographic characteristics of participants are presented in **Table S1**.

### Sample Collection and Processing

For the collection of endometrial samples, study participants reported that they were not actively trying to conceive and had not used hormonal treatment in the 3 months prior to recruitment. Once enrolled, study participants were instructed to use condoms during sexual intercourse for the duration of their enrollment. Endometrial biopsies were performed with the Endocell®, a disposable endometrial cell sampler (Wallach Surgical Devices, Trumbull, CT, USA) in the proliferative phase between cycle days 10-13 (n = 6) and during the early (n = 10), mid (n = 7), or late (n = 3) secretory phases, timed based on the urinary LH surge. On the day of the endometrial biopsy, blood (5 mL) was collected in nonheparinized, serum separator tubes, allowed to clot, and centrifuged at 4C. Serum estradiol (E2) and progesterone (P4) levels were determined (LabCorp, Raritan, NJ). Correct timing of biopsies was confirmed with serum E2 and P4 levels and endometrial histopathology as determined by two gynecological pathologists who were blinded to the timing of the tissue collection^13^.

For collection of endometrial deciduae, women scheduled for an elective termination of an uncomplicated pregnancy in the first trimester at 6-8 weeks estimated gestational age were eligible for inclusion. Study participants (n = 3) were recruited and enrolled prior to their scheduled surgical procedure. After dilation and curettage, tissue samples were floated in phosphate buffered saline (PBS, pH 7.2) and the endometrial decidua was isolated from the gestational sac and chorionic villi by mechanical separation using forceps.

After collection, endometrial tissue was placed in a tissue culture dish and rinsed with ice-cold PBS to remove blood and mucus. Using fine forceps, each sample was separated such that approximately 100 mg endometrial tissue was fixed in formalin and stored at room temperature for immunohistology; approximately 100 mg was preserved in RNAprotect (Qiagen, #76160) and stored at -80°C for RNA extraction for bulk RNA sequencing; and the remaining tissue was placed on ice, minced and processed within one hour of collection for scRNA-seq.

### Isolation of Endometrial and Decidual Cells

Cells were isolated from minced endometrial tissues for scRNA-seq by incubation at 37°C in digestion media (DMEM/F12, 3% charcoal-stripped fetal bovine serum (FBS), 1 mg/mL collagenase A (Roche, #11088793001), 0.1 mg/mL DNAse type I (Roche, #10104159001) with agitation on a benchtop shaker at 250 rpm for 15 minutes. The tissue was then further dissociated by repeatedly passing it through a 16G needle attached to a 10 mL syringe. The shaking and needle passage steps were repeated once more to ensure thorough dissociation. The cell suspension was centrifuged at 300× 𝑔 for 5 minutes. The resulting pellet was resuspended in 1 mL of TrypLE Select Enzyme (Thermo Fisher #12563011) containing 0.1 mg/mL DNase I and incubated at 37°C on a shaker at 250 rpm for 15 minutes. 10 mL of ice-cold digestion media was added, and the cell suspension was filtered through a 40 µm cell strainer. Red blood cells (RBC) were lysed using RBC lysis buffer (ThermoFisher, #00-4333-57) and dead cells were removed using the Dead Cell Removal Kit (Miltenyi Biotec, #130-090-101). The resulting single-cell suspension was used for Chromium (10X Genomics) sequencing.

### RNA Isolation, Library Preparation, and Sequencing

Total RNA was isolated from endometrial tissue using a RNeasy mini kit (Qiagen, #74136). RNA was quantified by Qubit (Invitrogen) and quality was assessed with a Fragment Analyzer (Advanced Analytical Technologies, Inc.) at Albert Einstein College of Medicine Epigenomics Shared Facility (RRID:SCR_023284). Libraries were prepared using the KAPA Stranded RNA-Seq Kit with RiboErase for Illumina Platforms (Kapa Biosystems #KK8483) with the addition of Ambion External RNA Controls Consortium (ERCC) spike-in controls (Invitrogen #4456740). Libraries were quantified, multiplexed, and sequenced as single end (1 x 75 bp) on an Illumina NextSeq 500 instrument (RRID:SCR_014983) to yield approximately 50 million reads per sample. FASTQ files were generated using picard (v2.26.10, RRID:SCR_006525) module ExtractIlluminaBarcodes followed by IlluminaBasecallsToFastq with default parameters, except “INCLUDE_NON_PF_READS = false”. Raw FASTQ files were trimmed of flanking adapter sequences using Trim Galore (v0.6.7, RRID:SCR_011847)^87^ with default parameters and “adapter = AGATCGGAAGAGC”. Read quality was assessed using FastQC (v0.11.9, RRID:SCR_014583) and FastQ Screen (v0.6.5, RRID:SCR_000141)^88^. Raw data were deposited at the NCBI Gene Expression Omnibus (GSE289073).

For scRNA-seq, libraries were generated and sequenced using the 10X Chromium Single Cell 3ʹ GEM kit (10X Genomics, v2). Paired end, 2 x 75 bp, sequencing was performed on an Illumina NextSeq 500 instrument (RRID:SCR_014983). Output was demultiplexed and converted to FASTQ format with Cell Ranger (10X Genomics, v3.1.0). Raw data were deposited at the NCBI Gene Expression Omnibus (GSE290822).

### RNA Sequencing Analysis

Trimmed reads were mapped to Homo sapiens genome assembly GRCh38 (hg38) using STAR (v2.7.9a, RRID:SCR_004463)^89^. Reads overlapping Ensembl^90^ annotations (v110) were quantified with STAR prior to model-based differential expression analysis using the edgeR-robust method^91–93^. A read counts matrix was used as input for the endest^94^ R package (v0.1.1) results were visualized with ggplot2 (v3.5.1, RRID:SCR_014601). For differential expression analysis, genes with low counts per million (CPM) were removed using the filterByExpr function from edgeR (RRID:SCR_012802). Genes were considered differentially expressed if the FDR-corrected p-values were less than 0.05. Venn diagrams were generated with the R package eulerr (v7.0.2, RRID:SCR_022753).

For scRNA-seq, demultiplexed sequencing reads were processed and aligned to the *Homo sapiens* genome assembly GRCh38 (hg38) using STAR (v2.7.9a) with 10X Genomics Cell Ranger (v3.1.0, RRID:SCR_017344). Samples were merged using the integration anchors function of the Seurat package (v5.1.0, RRID:SCR_016341) in R^95^. Genes expressed in fewer than three cells in a sample were excluded, as well as cells that expressed fewer than 200 genes and mitochondrial gene content >5% of the total unique molecular identifier count. Data were normalized using a global-scaling normalization method^95^ that normalizes the feature expression measurements for each cell by the total expression, multiplies this by a scale factor (10,000), and then log-transforms the results. High *HLA-G* and *CGA* cells were identified in pregnant biopsies as trophoblast were removed from our analysis. The top 2,000 most variable genes that were used for cell clustering were found using the *FindVariableFeatures* function and were then normalized using the *ScaleData* function. Based on an elbow plot generated using the *Elbowplot* function of Seurat, we selected 20 principal components (PC) for downstream analyses. Cell clusters were generated using *FindNeighbors* and *FindClusters* functions. For visualization, UMAPs were generated using the *RunUMAP*, *FeaturePlot* and *DimPlot* functions. The *DotPlot* Seurat function was used to generate dot plots to visualize gene expression for each assigned cluster. The Seurat function *AddModuleScore* was used to calculate ERA and decidualization scores. Ligand-receptor cellular communication analysis was determined with the CellChat (v2.1.2, RRID:SCR_021946)^43,44^ R package. Briefly, communication probabilities were calculated between cell types using the functions identifyOverExpressedGenes and identifyOverExpressedInteractions. Signaling sources, influencers, targets, mediators, and high-order information were obtained by communication network analysis. The resulting interactions were visualized using the included plotting functions.

### Glandular Epithelium Receptivity Module (GERM) Score Calculation

The receptivity module signature genes (n = 556, Table S7) were separated by positive or negative fold change and described as “GERM_up” or “GERM_down”, respectively. The resulting dataframe was used as input for the fgsea^96^ enrichment analysis implementation in clusterProfiler (v4.12.6, RRID:SCR_016884)^97^ with multiple datasets. Microarray and bulk RNA-seq experiments were downloaded from NCBI GEO (RRID:SCR_005012), using the GEO2R tool, as a list of genes and fold changes for early versus mid-secretory endometrium. Spatial and scRNA-seq datasets were further subset for the glandular or secretory glandular epithelium cell types. A combined normalized enrichment score (NES) was calculated as 𝐺𝐸𝑅𝑀 𝑠𝑐𝑜𝑟𝑒 = 𝐺𝐸𝑅𝑀_𝑢𝑝 + −1(𝐺𝐸𝑅𝑀_𝑑𝑜𝑤𝑛) and adjusted p-values were combined using the metap package with the sumlog function. The results were compiled and then visualized as a dotplot with ggplot2 (v3.5.1, RRID:SCR_014601).

### Immunostaining and Imaging

Formalin-fixed and paraffin-embedded endometrial tissue was sectioned at 5μm, placed on charged glass slides (StatLab, Millenia 1000), air-dried and stored at room temperature before use. Sections were stained with hematoxylin and eosin (H&E) or immunohistochemistry (IHC) staining. For IHC, sections were deparaffinized followed by antigen retrieval with BOND Epitope Retrieval Solution 2 (Leica Biosystems, USA) for 40 minutes. Primary antibodies to detect the estrogen receptor alpha (clone 6F11, Leica

Biosystems) and progesterone receptor (clone 16, Leica Biosystems) were applied for 20 and 24 minutes, respectively. Bound primary antibodies were detected with BOND Polymer Refine Detection kit. Sections were washed with distilled water, counterstained with hematoxylin, dehydrated through graded alcohols, cleared in xylene, and mounted with synthetic permanent media.

## Authorship Contribution Statement

**Gregory W. Burns:** Conceptualization, Methodology, Validation, Formal analysis, Data curation, Writing – original draft, Writing – review & editing, Visualization

**Emmanuel N. Paul:** Conceptualization, Methodology, Validation, Formal analysis, Data curation, Writing – original draft, Writing – review & editing, Visualization

**Manisha Persaud:** Conceptualization, Investigation, Writing – original draft, Visualization

**Qingshi Zhao:** Conceptualization, Methodology, Investigation, Writing – original draft

**Rong Li:** Conceptualization, Writing – original draft, Writing – review & editing

**Kristin Blackledge:** Investigation, Writing – original draft

**Jessica Garcia de Paredes:** Investigation, Writing – original draft

**Pratibha Shukla:** Investigation, Writing – original draft

**Ripla Arora:** Conceptualization, Writing – review & editing

**Anat Chemerinski:** Conceptualization, Formal analysis, Data curation, Writing – original draft, Writing – review & editing, Visualization, Supervision

**Nataki C. Douglas:** Conceptualization, Methodology, Validation, Formal analysis, data curation, Writing – original draft, Writing – review & editing, Visualization, Supervision, Project administration, Funding acquisition

## Acknowledgements

The authors would like to acknowledge Shahina B Maqbool, PhD, from the Epigenomics Shared Facility of the Albert Einstein College of Medicine and Robert Dubin, PhD, from the Epigenomics/Computational Genomics Core of the Albert Einstein College of Medicine Center. These studies have been funded by grants from the Eunice Kennedy Shriver National Institute of Child Health and Human Development K99HD112539 and SRI/Bayer discovery innovation grant to ENP, NIH R01HD109152 to RA and NIH R01AI148695 to NCD.

## Materials and Correspondence

Please address all correspondence to Nataki C Douglas.

