## Supplemental Figures for "Single cell mapping of the human endometrium and first trimester decidua identifies distinct epithelial and stromal cell contributions to fertility"

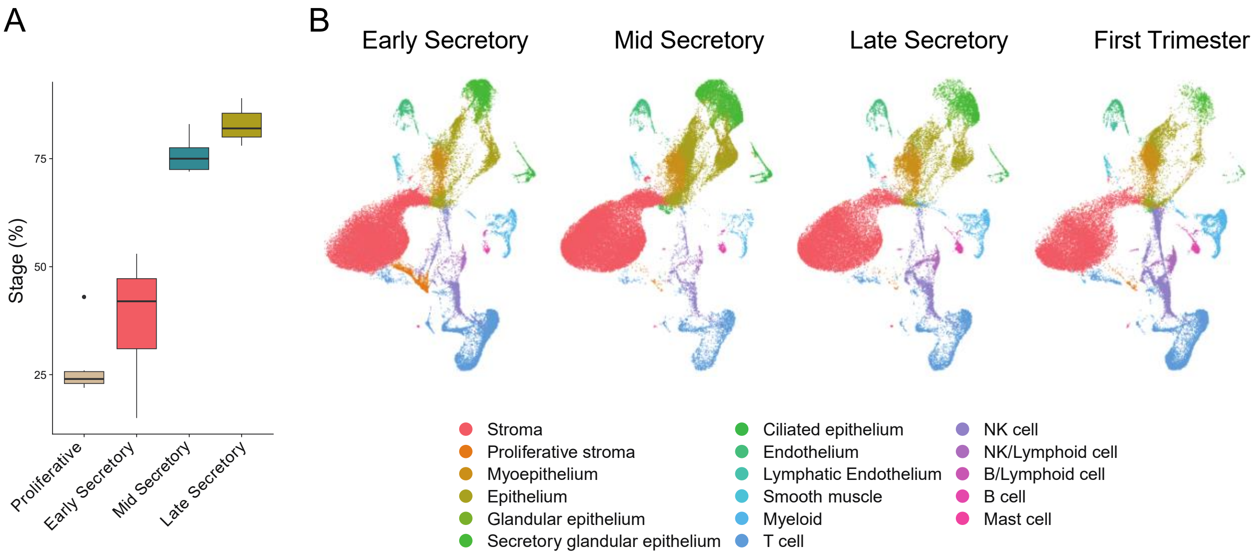


**Figure S1: Identification of the transcriptomic profile and dynamic cell populations in the endometrium over time.** (A) Boxplot of sample staging from the bulk RNA-seq using the method from Teh and colleagues[33]. (B) UMAP visualization of scRNA-seq data from human endometrial samples from the early (n = 4), mid- (n = 4), and late secretory (n = 3) phases, and first trimester decidua samples (n = 3).

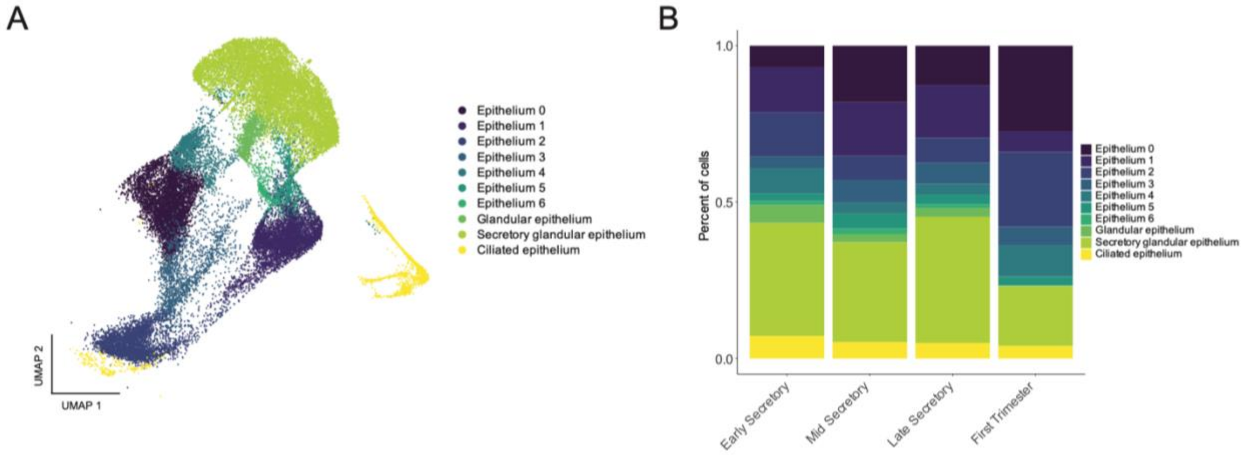


**Figure S2: Epithelium sub-clustering and dynamic distribution across stages.** UMAP visualization of scRNA-seq data from human epithelium sub-clusters (A). Stacked bar plot showing the proportion of cells in the epithelium subclusters by stage (B).

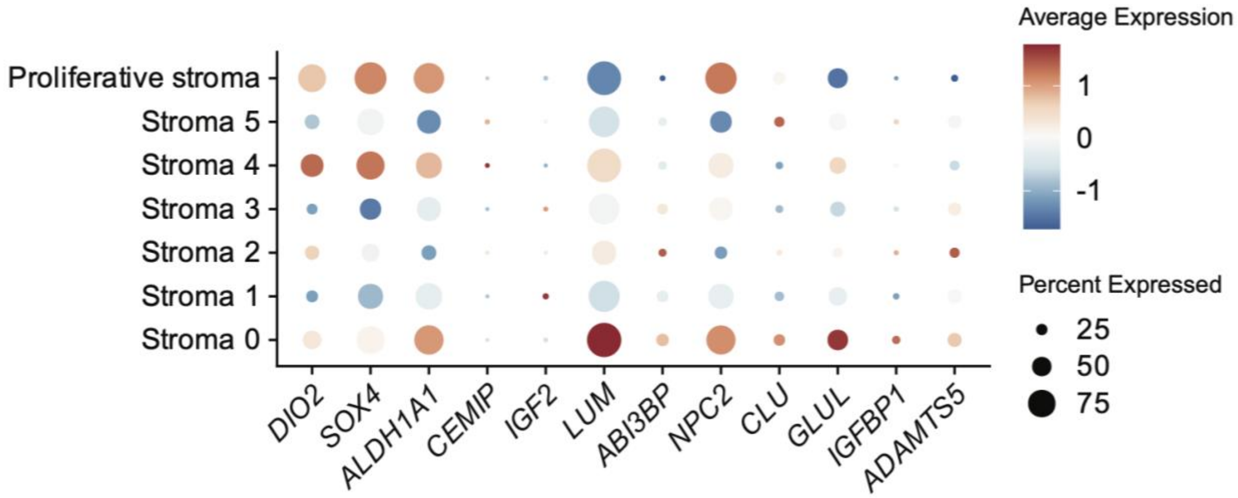


**Figure S3: Expression of i*n vitro* senescence markers is not restricted to a specific stroma subcluster.** Dotplot of *in vitro* senescence markers in the stroma subclusters.
