## Supplementary material for "Single cell mapping of the human endometrium and first trimester decidua identifies distinct epithelial and stromal cell contributions to fertility": Table S1

| Group | Subject | Age | Race | Ethnicity | BMI (kg/m^2) | Gravida | Para | E2 (pg/mL) | P4 (ng/mL) | Histologic dating | RNAseq |
|---|---|---|---|---|---|---|---|---|---|---|---|
| Mid-secretory | EMB 059 | 28 | Black or African American | Not Hispanic or Latino | 27.4 | 3 | 2 | 176 | 0.3 | PRO | Bulk |
| Mid-secretory | EMB 065 | 26 | Black or African American | Not Hispanic or Latino | 27.7 | 4 | 2 | 249 | 0.8 | PROFOC | Bulk |
| Mid-secretory | EMB 067 | 27 | Black or African American | Not Hispanic or Latino | 37.3 | 3 | 2 | 272 | 0.2 | WEAKLY PRO/SECRETORY | Bulk |
| Mid-secretory | EMB 068 | 29 | White | Hispanic or Latino | 35.5 | 3 | 3 | 82 | 0.2 | PROFOC | Bulk |
| Mid-secretory | EMB 070 | 23 | Black or African American | | 24.2 | 1 | 1 | 50 | 0.4 | PRO (GLANDS) | Bulk |
| Mid-secretory | EMB 077 | 40 | Black or African American | Not Hispanic or Latino | 47.8 | 4 | 3 | 195 | 0.8 | | Bulk |
| Early secretory | EMB 020 | 21 | Black or African American | Not Hispanic or Latino | 21.5 | 1 | 1 | 213 | 2.3 | PROFOC | Bulk |
| Early secretory | EMB 027 | 31 | Black or African American | Not Hispanic or Latino | 28.6 | 3 | 3 | 314 | 0.2 | PRO | Bulk |
| Early secretory | EMB 028 | 33 | Black or African American | Not Hispanic or Latino | 38.6 | 2 | 2 | 105 | 1.4 | PRO | Bulk |
| Early secretory | EMB 029 | 27 | Black or African American | Not Hispanic or Latino | 28.4 | 6 | 2 | 182 | 0.1 | PRO | Bulk |
| Early secretory | EMB 035 | 28 | Black or African American | Not Hispanic or Latino | | 2 | 2 | 168 | 0.4 | PRO | Bulk |
| Early secretory | EMB 037 | 32 | Black or African American | Not Hispanic or Latino | 49.2 | 4 | 4 | 158 | 0.5 | PRO | Bulk |
| Early secretory | EMB 044 | 30 | Black or African American | Not Hispanic or Latino | 38.2 | 1 | 1 | 191 | 0.8 | PRO | Bulk and single-cell |
| Early secretory | EMB 050 | 37 | Black or African American | Not Hispanic or Latino | 28.1 | 8 | 7 | 63 | 2.1 | PRO | Bulk and single-cell |
| Early secretory | EMB 051 | 37 | Black or African American | | 28.1 | 8 | 7 | 325 | 1 | PRO | Bulk and single-cell |
| Early secretory | EMB 053 | 23 | Black or African American | Not Hispanic or Latino | 26 | 2 | 1 | 106 | 1.2 | PRO | Bulk and single-cell |
| Mid-secretory | EMB 006 | 36 | White | Hispanic or Latino | 33.3 | 4 | 2 | 86 | 15.6 | 19-20D | Bulk |
| Mid-secretory | EMB 013 | 26 | Black or African American | Not Hispanic or Latino | 19.9 | 1 | 1 | 156 | 13.2 | 21-22D | Bulk |
| Mid-secretory | EMB 022 | 28 | Black or African American | Not Hispanic or Latino | 28.8 | 1 | 0 | 220 | 6.5 | 23-24D | Bulk |
| Mid-secretory | EMB 038 | 29 | Black or African American | Not Hispanic or Latino | 43.1 | 4 | 2 | 442 | 5.7 | 23-24D | Bulk |
| Mid-secretory | EMB 052 | 33 | Black or African American | Not Hispanic or Latino | 48.9 | 4 | 4 | 101 | 15.2 | 22D | Bulk and single-cell |
| Mid-secretory | EMB 055 | 24 | Black or African American | Not Hispanic or Latino | 26.8 | 3 | 3 | 150 | 12.9 | 22-23D | Bulk and single-cell |
| Mid-secretory | EMB 057 | 38 | White | Hispanic or Latino | 29 | 3 | 3 | 150 | 12.9 | 22-23D | Bulk and single-cell |
| Mid-secretory | EMB 087 | 41 | Black or African American | Not Hispanic or Latino | 33.5 | 5 | 5 | 76.5 | 11.4 | | single-cell |
| | EMB 066 | 32 | Black or African American | Not Hispanic or Latino | 29.8 | 3 | 2 | 131 | 4.8 | | Bulk and single-cell |
| | EMB 075 | 28 | White | Hispanic or Latino | 22.5 | 3 | 3 | 124 | 6.4 | | Bulk and single-cell |
| | EMB 088 | 31 | Black or African American | Not Hispanic or Latino | 39.8 | 6 | 2 | 107 | 4.8 | | Bulk and single-cell |
| | D 076 | 29 | Black or African American | Not Hispanic or Latino | 31.6 | 2 | 7 | | | | Bulk and single-cell |
| | D 079 | 24 | Black or African American | Not Hispanic or Latino | 34.2 | 4 | 1 | | | | Bulk and single-cell |
| | D 092 | 34 | Black or African American | Not Hispanic or Latino | 48.2 | 9 | 6 | | | | Bulk and single-cell |
